# Single-base mapping of m^6^A by an antibody-independent method

**DOI:** 10.1101/575555

**Authors:** Zhang Zhang, Li-Qian Chen, Yu-Li Zhao, Cai-Guang Yang, Ian A Roundtree, Zijie Zhang, Jian Ren, Wei Xie, Chuan He, Guan-Zheng Luo

**Affiliations:** MOE Key Laboratory of Gene Function and Regulation, State Key Laboratory of Biocontrol, School of Life Sciences, Sun Yat-sen University, Guangzhou, Guangdong, China; Laboratory of Chemical Biology, State Key Laboratory of Drug Research, Shanghai Institute of Materia Medica, Chinese Academy of Sciences, Shanghai, China; Department of Chemistry, Department of Biochemistry and Molecular Biology, Institute for Biophysical Dynamics, Howard Hughes Medical Institute, The University of Chicago, Chicago, IL, USA

**Author notes:** Correspondence should be addressed to G.Z.L.

## Abstract

*N*^6^-methyladenosine (m^6^A) is one of the most abundant mRNA modifications in eukaryotes, involved in various pivotal processes of RNA metabolism. The most popular high-throughput m^6^A identification method depends on the anti-m^6^A antibody but suffers from poor reproducibility and limited resolution. Exact location information is of great value for understanding the dynamics, machinery and functions of m^6^A. Here we developed a precise and high-throughput antibody-independent m^6^A identification method based on the m^6^A-sensitive RNA endoribonuclease (m^6^A-sensitive RNA-Endoribonuclease-Facilitated sequencing or m^6^A-REF-seq). Whole-transcriptomic, single-base m^6^A maps generated by m^6^A-REF-seq quantitatively displayed an explicit distribution pattern with enrichment on stop codons. Independent methods were used to validate the methylation status and abundance of individual m^6^A sites, confirming the high reliability and accuracy of m^6^A-REF-seq. We applied this method on five tissues from human, mouse and rat, showing that m^6^A sites were conserved with single nucleotide specificity and tend to cluster among species.

## Introduction

Among more than 100 RNA chemical modifications, *N*^6^-methyladenosine (m^6^A) is one of the most abundant form in eukaryotic mRNA, accounting for 0.1%-0.4% of all adenosine (*1*). The m^6^A modification influences various stages of mRNA metabolism as well as the biogenesis of lncRNA and microRNA, with affects across diverse biological processes including neuronal development, cell fate transition, immune response, DNA damage response, and tumorigenesis (*2–10*). Most m^6^A sites were found in conserved motif DRACH (D=G/A/U, R=G/A, H=A/T/C) (*1*), with partial methylation ratio ranging from 6% to 80% (*11*). Previous whole-transcriptome m^6^A maps have suggested that m^6^A modifications are enriched around stop codon, implying the functional importance of distribution pattern for m^6^A (*12–14*).

The chemical properties of m^6^A is similar to adenosine, making it difficult to discriminate by chemical reactions (*15*). Recently developed high-sensitivity mass spectrometry (LC-MS/MS) and blotting methods relying on antibodies were widely used to quantify the overall m^6^A level. To assess individual sites, methyl-sensitive ligase has been applied to confirm the methylation status of specific adenosines (*16, 17*), while the method called SCARLET can quantify the methylation level of individual m^6^A site (*11*). Other methods were developed to identify m^6^A in single-base resolution during reverse transcription, taking advantage of m^6^A-sensitive reverse transcriptase (*18, 19*), chemoenzymatic substitution of the *N*^6^-methyl group (*20*) or selective dNTP analogue such as 4SedTTP (*21*). Along with the rapid progress of single-molecule sequencing technologies, ONT sequencing platform is capable to detect modifications on model RNA (*22*), though the systematic error prevents the practical application in biological samples. Nevertheless, comprehensive interrogating of m^6^A at the transcriptome level is pivotal to reveal the biological importance of this mRNA modification. Methylated RNA immunoprecipitation and sequencing (MeRIP-seq or m^6^A-seq) has been widely used to profile m^6^A, identifying approximate region with m^6^A in ~100 nt length, while the exact location of individual m^6^A site remains undetermined (*12, 13*). Many refined methods have been developed to improve the resolution, such as PA-m^6^A-seq, miCLIP and m^6^A-CLIP (*23–26*). However, all of these methods are dependent on m^6^A-specific antibodies, suffering from poor reproducibility and complicated process. In addition, affinity variation and batch effects of antibodies make it difficult to quantify the methylation level (*27*). Therefore, a convenient and single-base resolution method is still needed for whole-transcriptome m^6^A identification and quantification, advancing the comprehension of m^6^A for its dynamics and cellular functions in post-transcriptomic regulation.

Several DNA endonucleases belonging to the restriction-modification (R-M) system have demonstrated sensitivity to DNA methylation. For instance, DpnI specifically cleaves methylated sites while DpnII is blocked by the same modified sequence. This feature has been adopted in genome-wide studies to detect the DNA 6mA modification in single base resolution (6mA-RE-seq/DA-6mA-seq) (*28*). For RNA endoribonucleases, a large group of sequence-specific enzymes belonging to the bacterial type II toxin-antitoxin (TA) system have been uncovered and the cleavage motifs are determined. Recently, an *Escherichia coli* toxin and RNA endoribonuclease, MazF, was reported to be sensitive to m^6^A modification within ACA motif, specifically cleaving the unmethylated ACA motif, leaving methylated (m^6^A)CA motifs intact (*29*). By screening the endoribonuclease pool, we identified endoribonuclease ChpBK as an endoribonuclease which can discriminate m^6^A-modified motifs from unmodified RNA. Taking advantage of the m^6^A-sensitive endoribonucleases, we developed m^6^A-sensitive RNA-Endoribonuclease-Facilitated sequencing method, m^6^A-REF-seq, which can identify transcriptomic m^6^A sites and quantify the methylation level in single-base resolution. To validate the m^6^A sites identified by this method, we employed a ligation-based method testing individual sites and achieved recurrent results independently. Further analysis revealed the hidden distribution pattern of m^6^A in single nucleotide resolution. Finally, we applied m^6^A-REF-seq to five tissues from human, mouse and rat, revealing the conservation of m^6^A in both single-base and regional levels across diverse tissues and species.

## Results

### Discrimination and quantification of m^6^A by endoribonuclease

The application of methylation sensitive DNA endonuclease in genome-wide 6mA identification inspired us to inspect the possibility of finding similar tools for m^6^A determination (*28*). To discover endoribonuclease with m^6^A sensitivity, we expressed and screened the candidate enzymes by testing the cleavage capacity to synthetic RNA oligonucleotides with or without m^6^A. Two enzymes, MazF and ChpBK, both belonging to the bacterial toxin-antitoxin system, were able to distinguish *N*^6^-methyladenosine (m^6^A) from unmethylated A. MazF recognized and cleaved the motif sequence ACA from the 5’-side of first A, leaving the methylated (m^6^A)CA motif intact (Fig. 1A). ChpBK was also blocked by m^6^A within the recognition motif, UAC (Fig. 1B). Synthetic RNA oligos with different m^6^A/A ratios were used to simulate the partially methylated mRNA. The cleavage assay demonstrated the fraction of digested oligo versus intact part was proportional to the methylation ratio, implying MazF has the potential to quantify the methylation level of mRNA (Fig. 1, C and D and fig. S1).

**Fig. 1.**
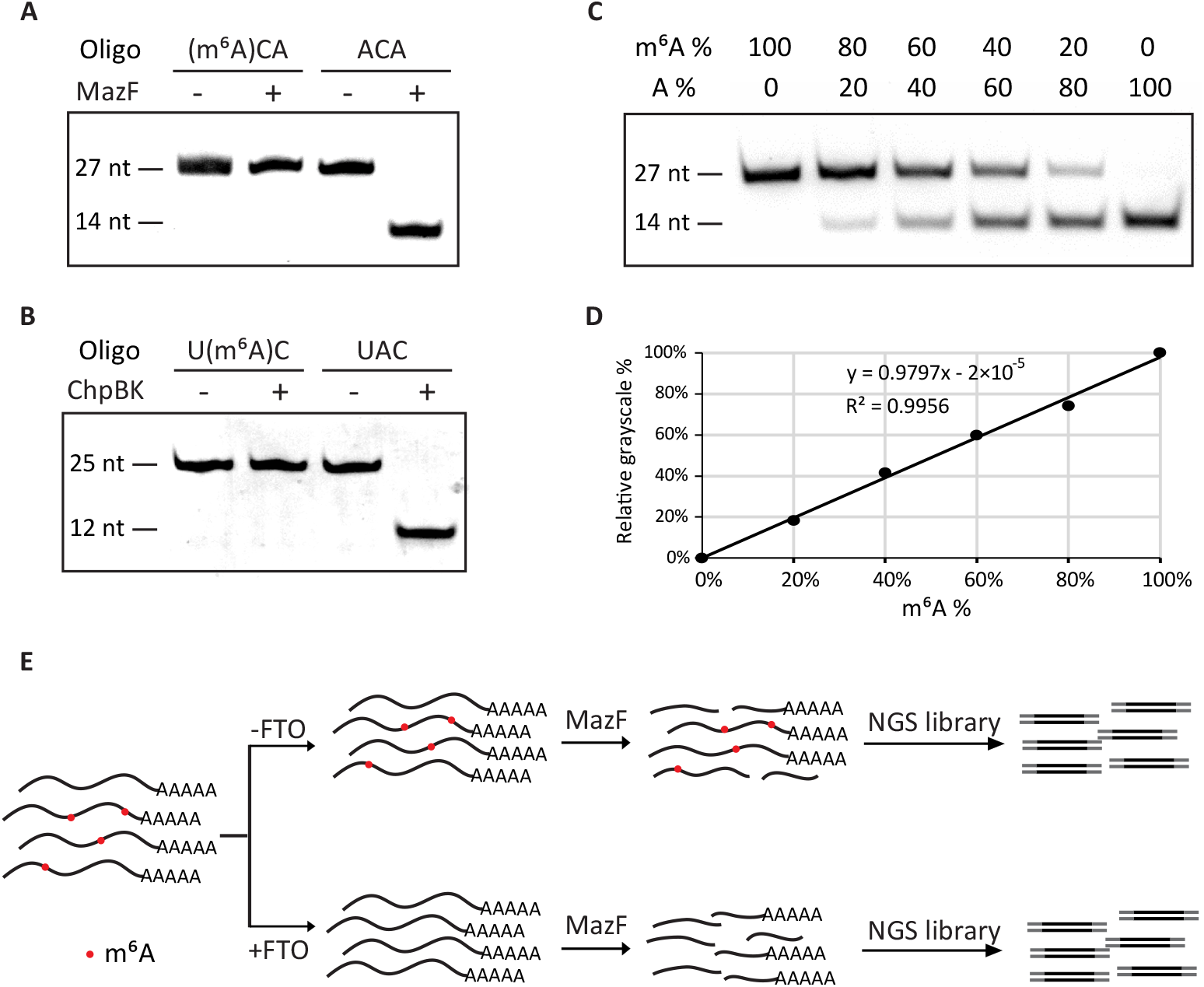
m^6^A identification method based on m^6^A sensitive RNA endoribonuclease. (**A**) Validation for the methylation sensitivity of endoribonuclease MazF by synthetic m^6^A-containing RNA oligonucleotide. (**B**) Validation for the methylation sensitivity of endoribonuclease ChpBK. (**C**) Various ratios of m^6^A-containing oligo mixed with unmethylated oligo digested by endoribonuclease. The m^6^A/A ratios of synthetic oligo are indicated to imitate practical RNAs with different m^6^A ratios. (**D**) Relative grayscale of proportionally mixed RNA oligo digested by endoribonuclease. (**E**) The schematic diagram of m^6^A-REF-seq.

The high sensitivity and specificity of enzymatically distinguishing m^6^A urged us to develop a new high-throughput method to identify single-base m^6^A on the transcriptome level. Since the MazF recognition motif ACA is more prevalent than that of ChpBK among known m^6^A sites, we chose MazF in following studies. Generally, mRNA from HEK293T cells was digested into fragments by MazF. After end-repair and purification, the mRNA fragments were ligated to NGS adaptors and amplified by PCR. To prevent potential false positives caused by unknown factors, a parallel sample batch demethylated by FTO in advance was also processed as the negative control (fig. S2). We call this method m^6^A-REF-seq, standing for m^6^A-sensitive RNA-Endoribonuclease-Facilitated sequencing (Fig. 1E).

### Transcriptome-wide identification of m^6^A sites in single-base resolution

We developed a pipeline to analyze the high-throughput sequencing data generated by m^6^A-REF-seq (Fig. 2A). As we anticipated, most reads contained the ACA at the 5’ terminus, in line with the effect of MazF treatment (fig. S3). The intact ACA motif sequence observed internally within an RNA fragment was supposed to be methylated. We verified this using the known m^6^A site with ACA motif on the 18S rRNA as an example. The unmethylated rRNA fragments were cleaved on the ACA motif and the methylated fragments were retained (Fig. 2B). Next we scanned each ACA motif within the transcriptome and interrogated the methylation sites by calculating the reads harboring intact internal ACA motifs. To further eliminate the potential false positive caused by RNA structure, we predicted the probability of each RNA fragment forming secondary structure according to minimum free energy (*30*), and removed the candidate sites tending to reside in double-stranded regions. We performed three biological replicates and retained the recurrent candidate sites detected in at least two replicates (Fig. 2C and table S1). For each site, the ratio of sequence reads with internal ACA motif versus reads split at the end represents the relative methylation ratio. We compared the methylation ratio of each candidate site in MazF treatment to the sample treated with FTO in advance, which was considered as demethylated RNA. Most of m^6^A sites had dramatic reductions in methylation level with FTO treatment (Fig. 2D). Only candidate sites with a significant decrease of methylation ratio in FTO treatment were considered as accurate m^6^A sites. In total we identified 4,260 high-confidence m^6^A sites for further studies.

**Fig. 2.**
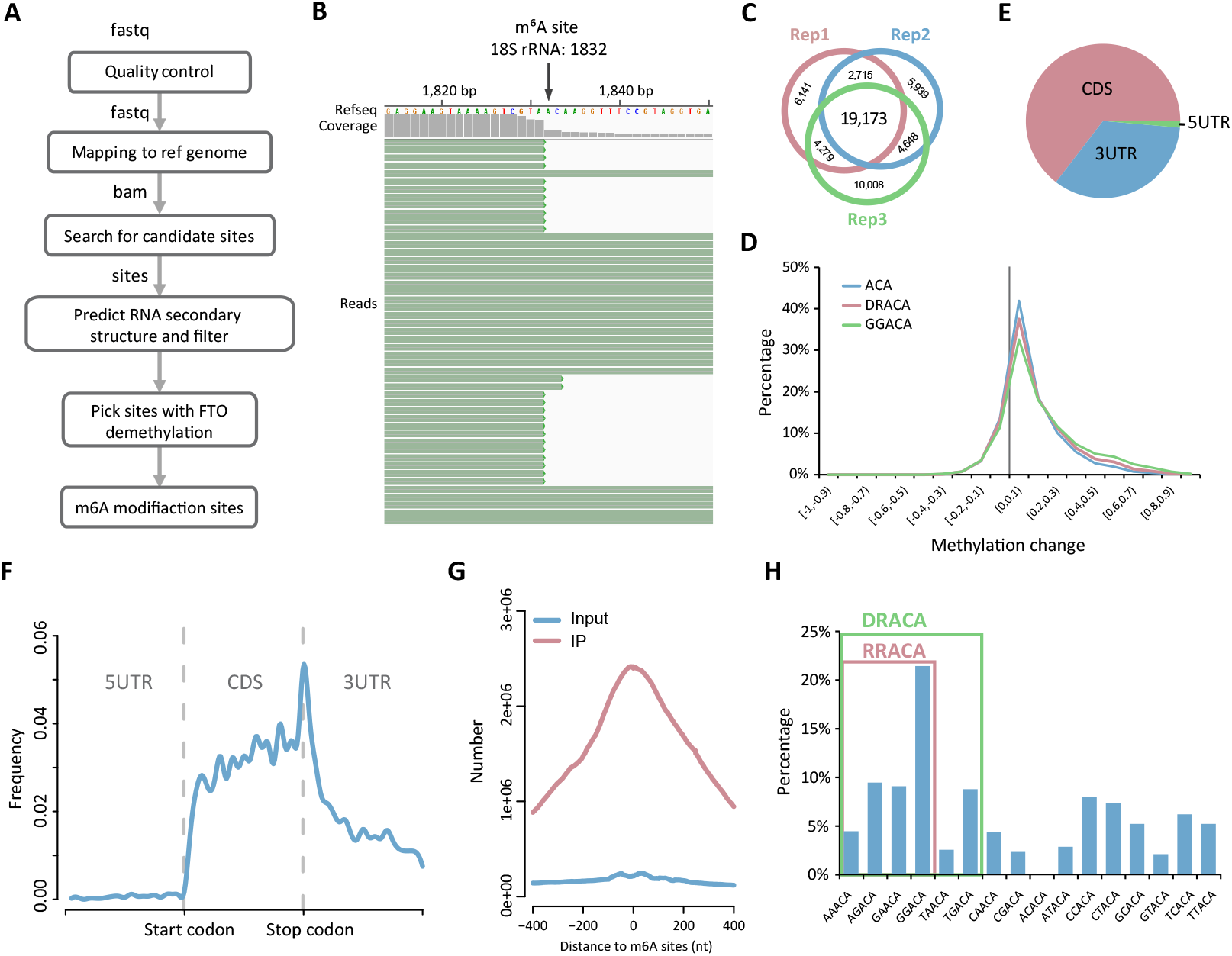
Transcriptome-wide distribution of m^6^A revealed by m^6^A-REF-seq. (**A**) The analysis pipeline of m^6^A-REF-seq data. (**B**) The snapshot of sequencing reads shows a known m^6^A site in 18S rRNA. (**C**) Overlap sites among three replicates after removing RNA secondary structure. (**D**) The overall shift of methylation ratio after FTO treatment. The value of methylation change indicates the methylation ratio of each m^6^A site in MazF minus that after FTO treatment. (**E**) Transcriptome-wide distribution of m^6^A. Pie chart shows the percentages of m^6^A sites located within CDS, 5’ UTR and 3’ UTR. (**F**) Single base m^6^A sites from m^6^A-REF-seq shows a typical transcriptome-wide distribution for m^6^A. (**G**) Overlap of m^6^A sites to the m^6^A peaks identified by antibody-based method. (**H**) The proportion of motifs containing m^6^A sites. The red square includes the RRACA motif, the green square includes the DRACA motif.

About 65% of these sites located in the CDS region and others were in UTRs (Fig. 2E). The transcriptome-wide distribution showed strong enrichment surrounding the stop codon (Fig. 2F), in line with the typical pattern discovered by antibody-dependent methods such as MeRIP-seq or m^6^A-seq. To compare the m^6^A location with MeRIP-seq in detail, we downloaded the profiling data deposited in GEO database (GSE29714) (*12*), and calculated the cumulated reads distribution of input and IP samples within the range of −400 nt to 400 nt flanking to every m^6^A site identified by m^6^A-REF-seq. The reads from IP samples were dramatically concentrated in this region, with a peak summit at almost the exact position of individual m^6^A sites; whereas reads from input sample did not show any enrichment, implying the m^6^A sites identified by m^6^A-REF-seq coincided well with the m^6^A peaks reported by MeRIP-seq (Fig. 2G). By expanding the context sequences flanking m^6^A sites, we found GGACA was the most enriched consensus motif. More generally, the consensus motifs DRACA and RRACA were overrepresented, accounting for 56% and 45% of all identified m^6^A sites, respectively (Fig. 2H). These results were in concordance with previous studies, in which DRACH/RRACH was the most preferred m^6^A motif in native mRNA (*12, 13, 24*).

### High reliability of individual m^6^A sites identified by m^6^A-REF-seq

To show the reliability of m^6^A-REF-seq as a new method, we sought another independent principle to validate individual m^6^A sites. The T3 ligase was reported to be sensitive against m^6^A sites during the ligation reaction, which had been applied practically to identify m^6^A sites in mRNA (*17*). We adopted this method and designed the probe L and probe R based on the sequence flanking the exact site (table S2), then used the T3 ligase to concatenate the two probes to an integrated template which could be amplified by PCR. According to previous result, the ligation efficiency at m^6^A site was significantly suppressed comparing to the efficiency at unmethylated A sites (*17*) (fig. S4). PCR amplification magnified the difference of ligation efficiency. Thus the amount of PCR products could be assessed on gel to represent the ligation efficiency, implying the methylation status of the interrogated site (Fig. 3A). *In vitro* FTO demethylation of mRNA was conducted, followed by the same ligase-based protocol for negative control. We picked nine m^6^A sites, including eight m^6^A sites and one unmethylated A. Six out of eight m^6^A sites were validated by the ligase-based method while the unmethylated site was also confirmed (Fig. 3B and fig. S5). To the best of our knowledge, this is the first batch of individual m^6^A sites confirmed by two independent methods. Interestingly, one validated m^6^A site (chr1: 29070177; Fig. 3B) is located at 343 nt upstream from the stop codon of m^6^A reader protein YTHDF2, suggesting a self-regulating feedback loop mediated by RNA modification.

**Fig. 3.**
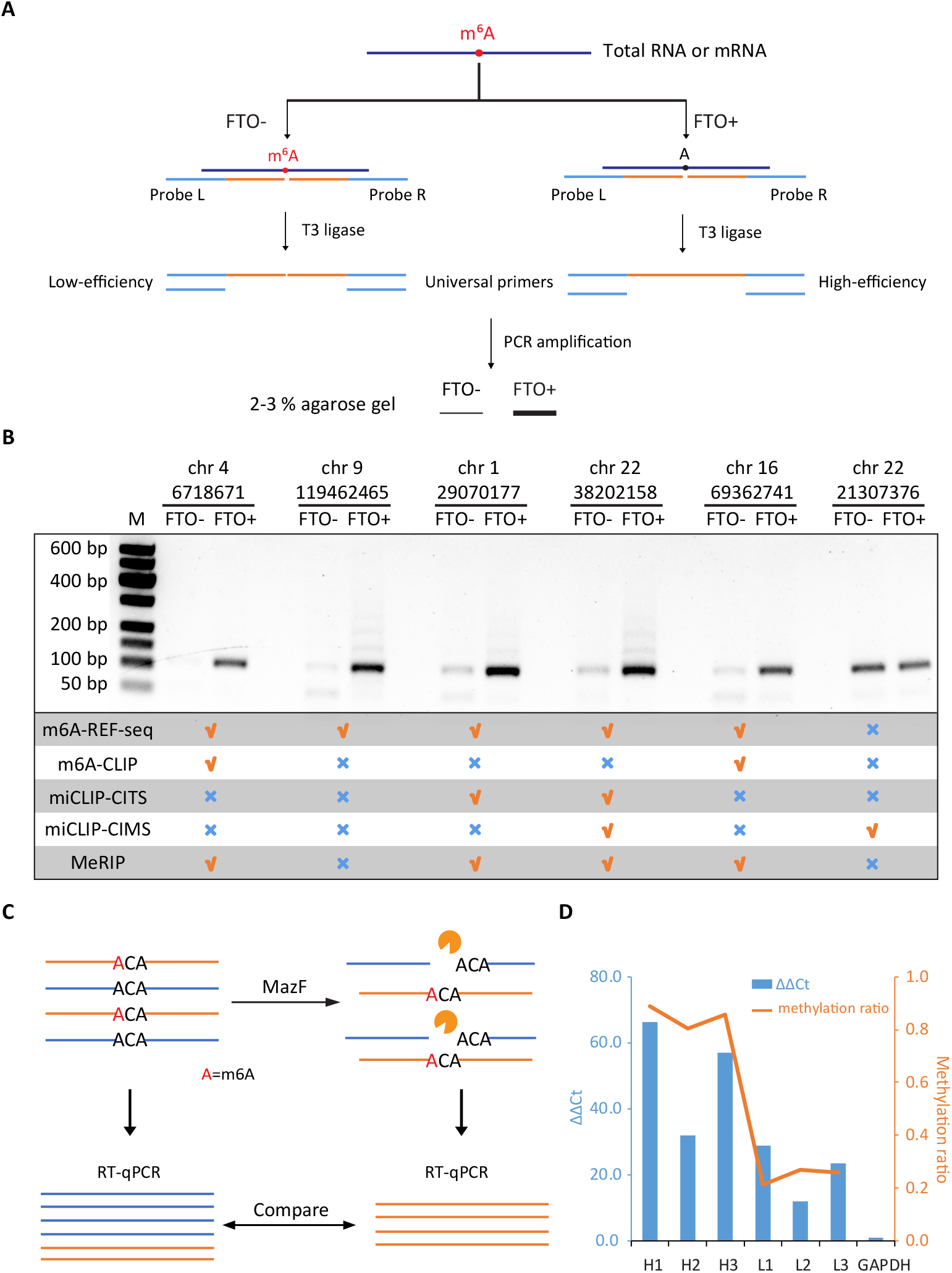
Single-base validation using ligation-amplification method and qPCR. (**A**) Schematic diagram of ligation-amplification method for single-base m^6^A validation. (**B**) Validation results of six individual sites. Five of the sites are validated to be m^6^A sites, whereas the other one is confirmed to be not modified. Data for m^6^A-CLIP, miCLIP-CITS, miCLIP-CIMS and MeRIP-seq are downloaded from published literatures. (**C**) Schematic diagram of qPCR to quantify the methylation level of a specific m^6^A site. The undigested mRNA sample is treated as control. (**D**) qPCR results and methylation ratios of six m^6^A sites. H1-H3 represent the highly methylated sites (> 0.75) while L1-L3 represent the weakly methylated sites (< 0.35). The left y-axis represents the ΔΔCt values and the right y-axis represents the methylation ratio determined by m^6^A-REF-seq.

Two other antibody-dependent methods, m^6^A-CLIP and miCLIP, had reported comprehensive m^6^A maps in transcriptome level. We checked whether the six individual m^6^A sites could be identified by these two methods. Peaks from MeRIP-seq data were also included as reference. Out of the six sites, one of them was not reported in either previous method. The negative control site supposed to be unmethylated was reported by miCLIP as an m^6^A site but not by other methods (Fig. 3B). We suspect that differences in samples, as well as the dynamic nature of m^6^A methylation contribute to these inconsistencies and potential false positives. Nevertheless, these results emphasize the need of independent methods to confirm authentic m^6^A sites for downstream analysis, especially considering m^6^A as a highly dynamic RNA modification.

### Quantification of m^6^A abundance by m^6^A-REF-seq

m^6^A-REF-seq not only determines the methylation status, but can also be used to quantify the methylation level of each m^6^A site by calculating the ratio of sequence reads with internal ACA versus reads split at the motif. To verify this, we designed a digestion-quantification assay that utilized RT-qPCR to evaluate the methylation level of identified m^6^A sites. First, mRNA was treated by MazF. The unmethylated sites located in ACA motif were cleaved, leaving intact methylated sites. After reverse transcription, cDNA spanning the intended m^6^A site was amplified by PCR to estimate the relative abundance. The same amount of mRNA without MazF digestion was amplified simultaneously as a control to compare with the digestion sample (Fig. 3C). We picked six m^6^A sites including three sites with high methylation level and three sites with low methylation level determined by m^6^A-REF-seq (table S3). The ΔΔCt values calculated from RT-qPCR results were comparable to the average methylation ratio determined by m^6^A-REF-seq (Fig. 3D and fig. S6), demonstrating m^6^A-REF-seq can principally quantify the methylation fraction of m^6^A sites.

### Characterizing m^6^A distribution pattern in single-nucleotide resolution

Previous studies demonstrated that one of most remarkable features of m^6^A was the distribution enrichment surrounding stop codon (*12, 13*). However, limited by the resolution of antibody-based methods, it was hard to know the precise position relative to the exact stop codon site, which divides coding regions and 3’ UTRs. The single-nucleotide m^6^A identification method largely increased the resolution of m^6^A map. Interestingly, our results showed this stop-codon-concentrated pattern could be reproduced by using single-nucleotide m^6^A sites, without any preference to coding regions or 3’ UTRs (Fig. 4A). Meanwhile, it was worth noting that consensus motif DRACA and DRACH were also enriched near stop codon, suggesting sequence motif might be the dominant factor to shape this pattern (Fig. 4A).

**Fig. 4.**
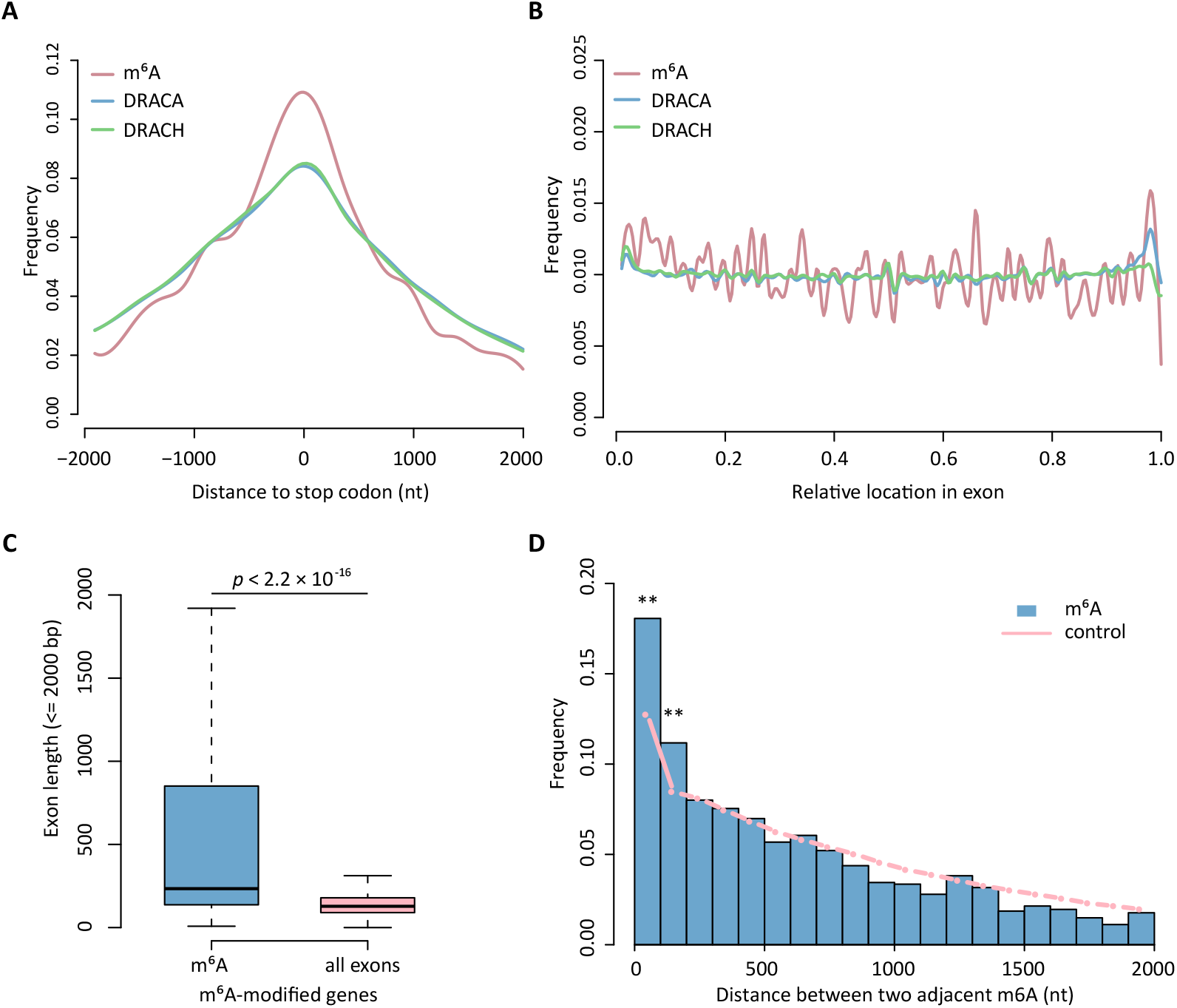
Single-base method reveals high resolution features of m^6^A. (**A**) The distance of individual m^6^A sites to the stop codons. DRACA and DRACH background motifs are extracted from the transcriptome. (**B**) The relative position of m^6^A sites within exons. (**C**) The length distribution of m^6^A-containing exons versus all exons in m^6^A-modified genes. (**D**) The distances between two m^6^A sites are significantly enriched within 200 bp regions compared to random sampling (Fisher’s exact test, **p < 0.01).

m^6^A had been proposed to be involved in various key regulatory processes of RNA metabolisms, including splicing (*31*). We calculated the relative distances of m^6^A sites to intron-exon boundaries in each exon and plotted in Fig. 4B. The result showed m^6^A distributed evenly along exon regions, with slight depletion towards the 3’ end. By mapping single m^6^A sites to exons, we compared the lengths of exons containing m^6^A modifications versus all exons. The lengths of m^6^A-containing exons were significantly greater than all exons (Fig. 4C, p < 2.2×10^-16^, Wilcoxon test), in line with previous report that m^6^A modification usually occurred within long exons (*13*).

Some indirect evidence showed that m^6^A tended to locate in short regions, suggesting a specialized working model of m^6^A as clusters. However, antibody-based methods could barely divide multiple sites from one m^6^A peak to demonstrate clustering. Utilizing the single-nucleotide m^6^A positions identified by m^6^A-REF-seq, we calculated the distances between each two adjacent m^6^A sites and compared to the control dataset, which contained the same number of sites randomly sampled from transcriptome 1,000 times. Our result showed two adjacent m^6^A sites were statistically prone to aggregate within 200 nt regions (p=6.6×10^-5^ and p=0.0065, Fisher’ exact test), supporting the hypothesis that m^6^A functions within clusters of modification (Fig. 4D).

### Conservation of m^6^A sites in mammals

We next applied m^6^A-REF-seq to identify m^6^A modifications in different tissues of human, mouse and rat, and identified more than 3,500 individual m^6^A sites in each sample (tables S4 and S5). The transcriptomic distribution displayed similar patterns among tissues in three species (Fig. 5A and figs. S7-S9). To examine the conservation of m^6^A at the individual gene level, we first collected 13,117 one-to-one orthologs in three species. By mapping the m^6^A sites to these ortholog genes in each type of tissues, we identified 2,439 human genes, 3,596 mouse genes and 1,343 rat genes in brain, among which 571 of them were shared in three species. The numbers were 536 and 262 for kidney and liver, respectively (figs. S8 and S9). These common ortholog genes modified by m^6^A in all three species were significantly more than expected, indicating a well-conserved distribution of m^6^A in gene level (Fig. 5B and table S6). Similarly, previous studies have reported the count of shared m^6^A peaks between human and mouse were significantly higher than expected, implying the conservation and important functions of m^6^A modification (*13*).

**Fig. 5.**
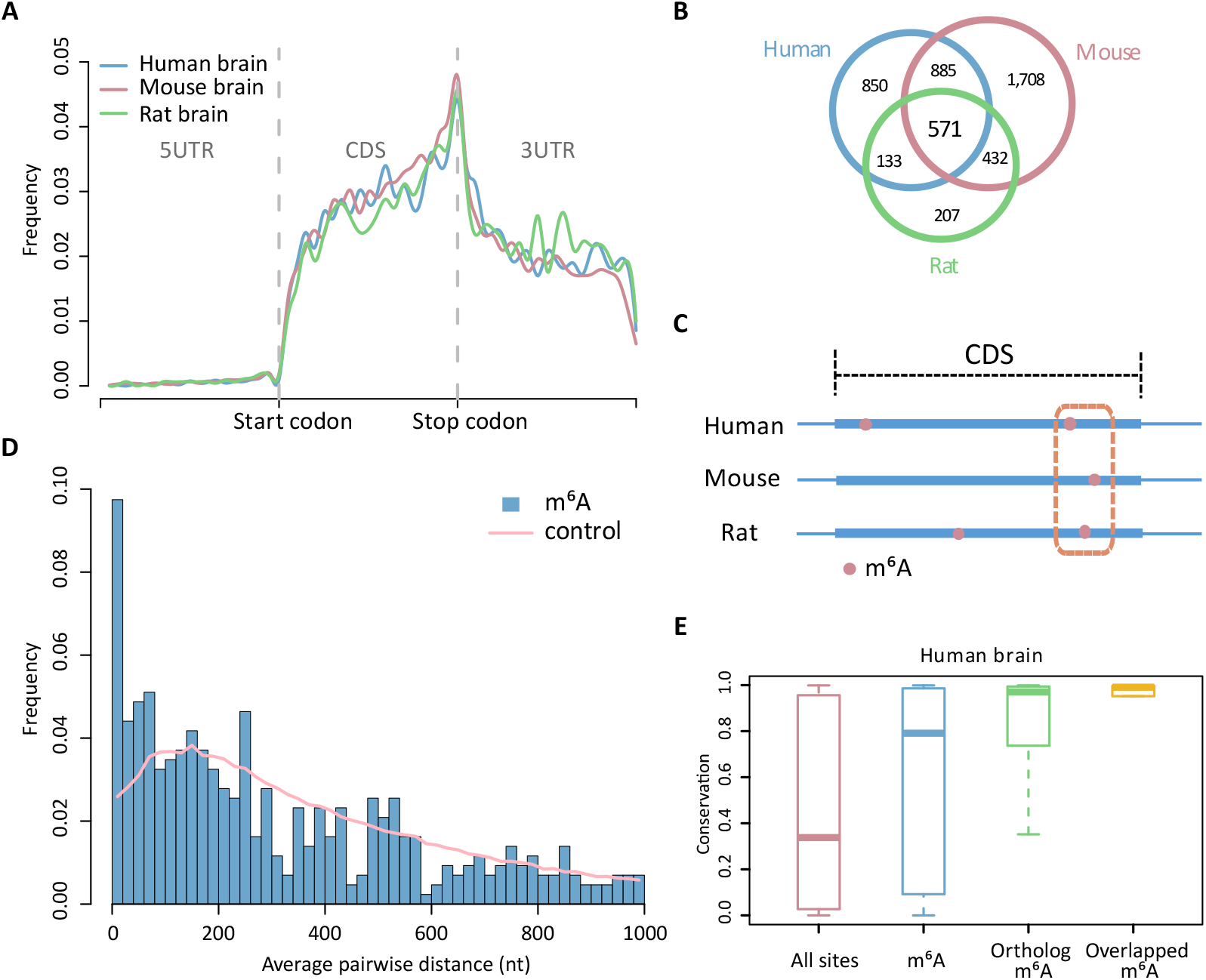
Conservation of m^6^A in mammals. (**A**) Metagene plots of m^6^A in the brain of human, mouse and rat. (**B**) Shared m^6^A-modified genes among three species. (**C**) Diagram showing the m^6^A sites conserved in the corresponding short regions from different species. (**D**) Frequency of distances for pairwise m^6^A in brain. Randomly picked ACA motifs are assigned for the same analysis as control. (**E**) Conservation scores of all m^6^A sites, methylation sites in ortholog genes and conserved m^6^A sites are compared to that for all A sites in ACA motifs (p-values < 9×10^-10^, Wilcoxon test).

Since m^6^A-REF-seq identified high confidence sites, we were able to investigate the conservation of m^6^A in single-nucleotide level. Another work analyzed the evolutionary conservation of yeast and human m^6^A sites in protein-coding regions and concluded that most m^6^A modifications were unconserved (*32*). However, the m^6^A sites used by this work were predicted based on the consensus motif DRACH, which could contain underestimated false-positives. Therefore, we mapped the single-nucleotide m^6^A sites identified by m^6^A-REF-seq to orthologous genes in three species, and calculated the significance of corresponding sites. In contrast to previous work, we found the counts of pairwise m^6^A sites were significantly more than expected (table S6). Furthermore, we tested whether the non-pairwise m^6^A sites tended to cluster in narrow corresponding regions among species. The shortest distances between pairwise m^6^A sites in ortholog genes were averaged (Fig. 5C). Random sampling of the same number of ACA motifs were performed for 1,000 times as control. The average distances between pairwise m^6^A sites were shorter than control (Fig. 5D, figs. S8 and S9), indicating that even though some m^6^A sites were not conserved in single-nucleotide level, they tended to cluster within conserved short regions across multiple species.

More extensively, we computed the PhastCons scores representing the conservation of each nucleotide among multiple-species (*33*). Identified m^6^A sites in brain were attributed and classified into sites conserved in three species or sites residing in ortholog genes. As expected, conserved m^6^A sites in single-base level and sites conserved in gene level had highest conservation scores (medians 0.991 and 0.970, respectively). The scores of overall m^6^A sites was also distinctively higher than that of all A sites in ACA motifs with the medians 0.791 and 0.338, respectively (Fig. 5E, p-values < 9×10^-10^, Wilcoxon test). Similar results hold for other tissues in human, mouse and rat (figs. S8-S10).

## Discussion

Antibody-dependent pulldown methods are widely used to profile transcriptomic m^6^A, the most prevalent modification in mRNA. An antibody-independent, easy-to-use method is strongly desired to cross-validate the known m^6^A sites and study the dynamic functions in singlenucleotide resolution. We developed m^6^A-REF-seq, an antibody-independent, high-throughput and single-base m^6^A detection method based on m^6^A-sensitive RNA endoribonuclease. High sensitivity and specificity of endoribonuclease we discovered to discriminate m^6^A modification, as well as stringent data process by removing potential RNA secondary structure and the introducing of FTO demethylation control, ensured reliability of m^6^A sites identified by this method. Transcriptome-wide m^6^A maps in human HEK293T cell and mammalian tissues displayed similar distribution pattern with previous antibody-based pulldown methods. To validate the m^6^A sites identified by m^6^A-REF-seq, we introduced another independent method which utilized T3 ligase to differentiate individual m^6^A or A at template RNA. The high consistency between different methods demonstrated the efficiency of m^6^A-REF-seq with high confidence. We also applied quantitative PCR (qPCR) to show that m^6^A-REF-seq could quantify the methylation level of individual m^6^A site.

The RNA endoribonuclease MazF has high sensitivity and specificity for RNA substrates. It works very well on extremely limited substrate, as little as nanograms or even picograms. Rapid and simplified experiment design, without the need for antibody-enrichment step largely reduces the requirement of starting RNA amount and sample preparation time. Therefore, this method could be applied to the studies for rare biological materials such as samples from pathological tissues or early embryos. Additionally, m^6^A-REF-seq offers the potential to capture the subtle changes of m^6^A during metabolic processes, advancing the dynamic studies of this post-transcriptional RNA modification in different life stages.

By screening RNA endoribonucleases, we discovered two m^6^A-sensitive enzymes from the bacterial type II toxin-antitoxin (TA) system, which recognized specific motifs “ACA” and “UAC”, respectively. To cover all the RRACH motifs in the transcriptome, we need new enzyme(s) capable of recognizing more universal sequence motifs while retaining the m^6^A-sensitivity. Based on our preliminary screening, we speculated that the m^6^A-sensitivity might be a general feature of endoribonucleases belonging to type II TA system. Further exploring in the endoribonucleases pool would be helpful to expend the toolbox for RNA modification studies. Another strategy is to transform the existing enzyme to adapt for new recognition sequences while retaining the methylation sensitivity. We studied the structure of MazF protein and mutated the 56th Lysine to Alanine. Even though the MazF-K56A mutant did not meet our expectation, the methyl-sensitivity was retained (fig. S11), suggesting single base or amino acid mutation may influence the cleavage efficiency or even change the target motif. On the other hand, directed evolution would be a promising way to expand the recognition sequence scope of MazF or other related proteins, as long as dedicated screening system was created.

We applied m^6^A-REF-seq to five different tissues from human, mouse and rat. The singlebase m^6^A maps displayed a distribution pattern similar to previous findings, that m^6^A tended to enrich surrounding stop codon. More importantly, this unique pattern seemed to be conserved among different species. However, because of limited resolution, previous methods could not tell to what extent the modification is conserved. In this study, we mapped the m^6^A sites to orthologous genes in these three species and compared the conservation score in single-nucleotide level. We found significantly more m^6^A-modified genes were shared among species than expected, and a large group of m^6^A sites could be aligned to the syntenic regions, showing the m^6^A modification was conserved at both the gene and individual base level. The well-conserved feature restated the biological importance of m^6^A, implying other uncovered roles of this RNA modification.

Taken together, m^6^A-REF-seq provides a new perspective for single-base m^6^A identification on the transcriptome level. The independent principle eliminates the need for m^6^A-specific antibodies, making it unique and amenable for future applications in small and precious samples. The features of this efficient and convenient method largely expand the scope of m^6^A studies.

## Materials and Methods

### RNA materials

The RNA oligonucleotides were synthesized from Takara Bio Inc as listed below.

RNA-m^6^ACA 5’ UUGGUUUUUUUUGG(**m^6^A**)CAUGUAUAUAGU 3’
RNA-ACA 5’ UUGGUUUUUUUUGGACAUGUAUAUAGU 3’
RB1-m^6^ACA 5’ GUUGUGUGAUAU(**m^6^A**)CAUAUGGUGGUG 3’
RB1-ACA 5’ GUUGUGUGAUAUACAUAUGGUGGUG 3’

Human HEK293T cell line was cultured with DMEM (Corning) supplemented with 10% (v/v) FBS (Gibco) at 37 □ in the presence of 5% CO_2_, and tested to be mycoplasma negative. Total RNA from HEK293T cells was extracted using Trizol reagent (Thermo Fisher Scientific) and the mRNA was purified using Dynabeads mRNA Purification Kit (Thermo Fisher Scientific, Cat. No. 61006). The mRNA of human tissues (brain, liver and kidney), mouse tissues (brain, liver, heart, testis and kidney) and rat tissues (brain, liver and kidney) were purchased from Takara Bio Inc.

### RNA endoribonuclease validation for MazF

RNA oligo (10 pmol) were incubated with 2.5 U MazF (mRNA intereferase-MazF, Takara, code No. 2415A) in the 20 ul reaction mixture of MazF buffer (40 mM sodium phosphate (pH 7.5) and 0.01% Tween 20) at 37 □ for 30 min. The RNA oligos were mixed with the m^6^A percentage of 0%, 20%, 40%, 60%, 80%, 100%, and the oligo mixtures were incubated with 2.5 U MazF in the 20 ul reaction mixtures. The samples were loaded into 15% urea-polyacrylamide gel and electrophoresed in 0.5× TBE buffer.

The FTO demethylation reaction was conducted in the reaction mixture contained 10 pmol m^6^A RNA oligo, 0-10 ug FTO demethylase, 283 μM of (NH_4_)_2_Fe(SO_4_)_2_·6H_2_O, 300 μM of α-KG, 2 mM of L-ascorbic acid, 20 U RNase inhibitor (Takara, code No. 2313A) and 50 mM of Tris·HCl buffer (pH 7.5). The reaction was stopped by 40 mM EDTA after 3h incubation at room temperature. The MazF cleavage reaction was performed to test efficiency of FTO demethylation.

### Dot-blot assay of FTO demethylation

m^6^A FTO demethylase with different concentration (0 ug, 0.25 ug and 2.5 ug) were used in the dot-blot assay to test the demethylation of RNA. The total RNA was incubated with FTO, 283 μM of (NH_4_)_2_Fe(SO_4_)_2_·6H_2_O, 300 μM of α-KG, 2 mM of L-ascorbic acid for 3 h at room temperature. After purified with RNA Clean & Concentrator-5 kit (ZYMO Research), the concentration of total RNA was determined by Qubit RNA HS Assay Kit (Thermo Fisher Scientific) and diluted to 10 ul volume with 500 ng. Spotting and thermal crosslinking were carried out on a metal base plate preheated to 70 □. After thermal cross-linking for 15 min, UV cross-linking was conducted twice with 2000 KJ. Then, 3% BSA was used to block at room temperature for 1 h. The anti-m^6^A antibody (Synaptic Systems, Cat. No. 202003) was diluted to 1:2000 and incubated with the nylon membrane for 2 h. The nylon membrane was washed for three times with 1% TBST, and then was incubated with the 1:5000 diluted anti-rabbit antibody for 1 h. The membrane was washed for five times with 1% TBST. After draining off the water, the chemiluminescence detection signal was performed using the ECL system. The membrane was washed twice with 1% TBST. After draining off the water, it was dyed with methylene blue solution for 8 min, and then washed with 1% TBST. The signal was detected by a white light system.

### Expression and purification of recombinant ChpBK protein

To express recombinant ChpBK protein (NCBI Reference Sequence: NP_418646.1), *E. coli* BL21(DE3) strain carrying PET-28a-chpBK OD600 plasmid was grown in LB medium supplemented with Kan 10 ug/ml at 37 □ and 200 rpm until the OD600 reach to 0.5. Isopropyl-β-D-thiogalactopyranoside (IPTG) was added to a final concentration of 0.4 mM and cell growth resumed for 2 h under the same conditions. Harvested cell pellets were lysed by sonication in 30 ml of buffer A (10 Mm BisTris, pH 7.0, 30 mM imidazole), and centrifuged for 20 min at 16,000 g and 4 □. The soluble fraction was incubated on ice for 30 min in the presence of DNase and RNase (10 g/ml; Thermo Fisher Scientific). The His-ChpBK fusion protein was purified from clarified lysates using a Ni-NTA beads (Invitrogen). Protein purity was determined by SDS-PAGE analysis and samples were quantified using the Bio-Rad protein assay and stored at −80 □.

### Expression and purification of MazF-K56A in *E. coli*

The 56th Lysine of MazF protein was mutated to Alanine. The protein sequence is shown below, where the underlined bold “A” indicates the mutated amino acid.

>MazF-K56A

MVSRYVPDMGDLIWVDFDPTKGSEQAGHRPAVVLSPFMYNNKTGMCLCVPCTTQS<u>**A**</u>GY PFEVVLSGQERDGVALADQVKSIAWRARGATKKGTVAPEELQLIKAKINVLIG

The plasmid DNA pCOLD-MazF was transformed into *E. coli* Rossetta strain and a single colony was picked by sterilized loop and inoculated to 50 mL LB (containing Ampicillin) in sterile culture flask. *E. coli* was cultured at 37 □ until OD600 reached 0.6-0.8. IPTG (1 mM) was added to induce protein expression and cells grew overnight at 18 □. Cells were harvested by centrifugation and suspended in 200 ml lysis buffer (50 mM Tris pH 8.0, 350 mM NaCl, 10 mM Imidazole). Cell pellets were lysed by sonication and the supernatant were collected by centrifugation and applied to Nickel affinity purification using HisTrap HP. The column was washed by lysis buffer to remove non-specific binding protein and target protein was eluted with linear gradient of elution buffer (50 mM Tris pH 8.0, 350 mM NaCl, 500 mM Imidazole). We concentrated the protein and changed protein storage buffer to Ion-exchange binding buffer (20 mM HEPES, 50 mM NaCl, pH 7.5) by HiTrap desalting column. TEV protease was used to remove the His-tag and target protein was loaded to HiTrap HP Q column. Target protein was eluted by linear gradient of Ion-exchange elution buffer (20 mM HEPES, 1 M NaCl, pH 7.5). The concentrated protein was analyzed by SDS-PAGE.

### RNA endoribonuclease validations for ChpBK and MazF-K56A

The digestion reaction of ChpBK was conducted in a 20 ul reaction mixture with 10 pmol RNA oligo (RB1), 1 ug ChpBK, 20 U RNase inhibitor and 10 mM Tris·HCl buffer (pH 7.5). The mixture was incubated at 37 □ for 30 min. The digestion reaction of ChpBK with different m^6^A percentage was conducted as described above. The MazF-K56A cleavage reaction was performed in a 20 ul reaction mixture with 10 pmol RNA oligo, 2 ug MazF-K56A and 1× MazF buffer (40 mM sodium phosphate (pH 7.5) and 0.01% Tween 20) at 37 □ for 30 min. All samples were loaded on the 15% urea-polyacrylamide gel and electrophoresed in 0.5× TBE buffer.

### Library construction for next-generation sequencing

mRNA was first heated at 85 □ for 5 min and chilled on ice for 2 min to denature RNA secondary structure. The FTO demethylation reaction was conducted before MazF reaction in 20 ul of reaction mixture containing 100-200 ng mRNA, 2.5 ug FTO demethylase, 283 μM of (NH_4_)_2_Fe(SO_4_)_2_·6H_2_O, 300 μM of α-KG, 2 mM of L-ascorbic acid, 20 U RNase inhibitor and 50 mM of Tris·HCl buffer (pH 7.5). After incubating at room temperature for 3h, the reaction was stopped by adding 40 mM EDTA. In order to prevent the influence of FTO buffer, the MazF treatment was supplemented with the same reaction mixture except FTO protein. The MazF and FTO-MazF samples were heated at 85 □ for 5 min and chilled on ice for 2 min, and then incubated with 10 U MazF in the MazF reaction buffer at 37 □ for 30 min. The fragmented mRNA was purified by RNA Clean & Concentrator-5 kit (ZYMO Research) and eluted into 20 ul RNase-free water. The eluted RNA fragment was end-repaired by the T4 polynucleotide kinase (T4PNK; Vazyme Biotech) in 50 ul reaction mixture with 1x T4PNK buffer, and incubated at 37 □ for 30 min. After purification by RNA Clean & Concentrator-5 kit (ZYMO Research), the concentration of fragmented mRNA was measured by Qubit RNA HS Assay Kit (Thermo Fisher Scientific). The NGS library was constructed by using VAHTS Small RNA Library Prep Kit (Vazyme Biotech, NR801) with 10 ng fragmented mRNA and amplified by PCR for 15 cycles. Libraries were loaded on the 8% urea-polyacrylamide gel and electrophoresed in 0.5× TBE buffer, and purified libraries were sequenced on the Illumina HiSeq X10 platform. Three replicates have been conducted for mRNA of HEK293T cell line. The same protocol was also conducted for mammalian tissues.

### NGS data analyses

First, quality control was performed on raw sequencing reads by the open software FastQC (http://www.bioinformatics.babraham.ac.uk/projects/fastqc/). Then adaptors were filtered by cutadapt (*34*) with at least 15 nt remaining length of paired-end reads. The clean reads were mapped to the reference genome (hg19) using Hisat2 (*35*) with default parameters except reporting only one result. IGV was used for data visualization (*36*). We scanned the motif ACA (and the reverse complementary motif TGT) on the reference genome and mapping result file, respectively, counting the number of ACA/TGT at the internal or terminal of a sequence read. As short sequence reads with lengths less than 15 nt were filtered, only ACA/TGT with the distance from the end of a read more than 15 nt was treated as “internal” of a read. Each ACA/TGT motif on the reference genome which has more than ten reads supporting the motif located internal of reads were treated as a candidate m^6^A site. The ratio of sequence reads with internal ACA versus reads split at the end represents the relative methylation ratio of each site. Candidate sites with repeated or continuous ACA motifs were removed.

As the RNA secondary structure could affect the reaction efficiency, we predicted the probability of each RNA fragment forming secondary structure, and removed the candidate sites tending to reside in double-stranded regions. The RNAfold program of ViennaRNA package (*30*) was used to predict intramolecular secondary structure of each read which supported the candidate sites. The condensed representation of the pair probabilities of each nucleotide was parsed according to the tutorials of ViennaRNA package. The pairing probability value of NNACA motif was calculated and candidate sites with high pairing probability were discarded. The FTO demethylation treatment was also treated as a negative control. Candidate sites with at least 10% methylation ratio decrease were remained as m^6^A sites. The metagene plot of m^6^A sites, which was described the relative location of each site on the 3’UTR, CDS, and 5’UTR of mRNA, was calculated and plotted as well. The previous m^6^A-CLIP and miCLIP datasets were downloaded from Ke et al. (*25*) and Linder et al. (*24*), respectively. The raw reads of MeRIP-seq datasets were downloaded from Meyer et al. (*12*) (GSE29714) and mapped to human (hg19) reference genome using Hisat2. The reads within the range of −400 nt to 400 nt flanking to every m^6^A site were plotted. As the sequencing data was single-end, we extended the reads to 200 nt to simulate the IP peak.

### Single-base validation using ligase-based method

Refined single-base T3 ligase-based method (*17*) was used to validate m^6^A sites identified by m^6^A-REF-seq. The FTO demethylation sample was used as negative control (i.e. the FTO+ treatment in Fig. 3A). The ligation reaction mixture A consisted of 20 nM probe L, 20 nM probe R, 1× T3 ligation buffer (NEB) and 300 ng total RNA with or without FTO treatment. The ligation reaction mixture B consisted of appropriate amount of T3 ligase (NEB) with ligation buffer. Mixture A was heated at 85 □ for 3 min and then incubated at 35 □ for 10 min, and then the ligation reaction mixture B was added to the final volume of 10 ul. This mixture was incubated at room temperature for ligation reaction for 8-10 min and chilled on ice immediately. The volume of 1ul ligation product was used to PCR amplified for 24 cycles in the 2× Taq mix (Vazyme Biotech). The samples were loaded into the 2-3% agarose gel and electrophoresed in 0.5× TAE buffer to detect the signal.

### qPCR validation for m^6^A quantification

The MazF treatment was performed in 10 ul volume with 500 ng mRNA and 10 U MazF at 37 □ for 30 min. The digested mRNA and non-digested mRNA were subjected to reverse transcription using 5×TransScript All-in-One First-Strand cDNA Synthesis SuperMix for qPCR (One-Step gDNA Removal; Transgen Biotech, AT341-02). TB Green Premix Ex Taq II (Tli RNaseH Plus, Takara, Code No. RR820A) was used on BIO-CFX96 Touch.

### Characterizing m^6^A distribution pattern in single-base resolution

Consensus motifs DRACA and DRACH were scanned on the reference genome, and the distances of A sites in the two motifs to stop codons were calculated and plotted as well. The relative position in exon for m^6^A, regular A sites in DRACA and DRACH were computed. In order to estimate the distance between two adjacent m^6^A sites, only genes with more than two m^6^A sites were analyzed. We randomly sampled the same number of ACA motifs as the number of observed m^6^A and repeated this step for 1,000 times as control. The distances between adjacent A sites in control were calculated. Fisher’s exact test was used to test the significance of first two bins. The conservation analysis was conducted using PhastCons scores of human hg19 genome downloaded from UCSC.

### Conservation of m^6^A in the same mammal tissues

m^6^A-REF-seq protocol was used to identify m^6^A sites in the different tissues of human (brain, liver and kidney), mouse (brain, liver, heart, testis and kidney) and rat (brain, liver and kidney). The one-to-one orthologous genes in pairwise species of human (GRCh37.p13), mouse (GRCm38.p6) and rat (Rnor_6.0) were downloaded from Ensembl (Ensembl genes 94). A total of 13,117 one-to-one orthologous genes among three species were retained for downstream analyses. Protein sequences of 13,117 orthologs in human, mouse and rat were aligned by Muscle (*37*) and then converted to corresponding coding sequence alignments using PAL2NAL (*38*). The shared m^6^A-modified genes and share m^6^A sites among species were calculated and tested for significances. The average pairwise distances among closely m^6^A sites in three species were calculated and compared with random control sample. The control sample was randomly picked for the same number of ACA motifs as the number of observed m^6^A for 1,000 times. Analyses of conservation score were conducted by using PhastCons scores of human (hg19), mouse (mm10) and rat (rn6) downloaded from UCSC.

## Author contributions

Z.Z. C.H. and G.Z.L. conceived the project; Z.Z., L.Q.C. and Y.L.Z. conducted the experiments; Z.Z. analyzed the NGS data; Z.S.Z., I.A.R., C.G.Y., J.R. and W.X. provided the supporting materials and ideas; Z.Z., L.Q.C., C.H. and G.Z.L. wrote the manuscript. All authors reviewed the results and approved the final version of the manuscript.

## Data and materials availability

Data were deposited in NCBI’s Gene Expression Omnibus (GEO) under accession number GSE125240.

