## Supplementary file for "Single-base mapping of m^6^A by an antibody-independent method"

#### This file includes:

Supplementary tables S1-S6

Supplementary figures S1-S11

### Supplementary Tables

**Table S1. Basic information of sequencing data for human HEK 293T cell line.**

| Replicates | Treatment | Clean data (million reads) | Mapped reads (million reads) | Mean coverage (Q20*) | # covered transcripts |
|---|---|---|---|---|---|
| Rep1-MazF | MazF | 20.14 | 15.15 | 2.96 | 30,235 |
| Rep1-FTO | FTO-MazF | 33.29 | 24.90 | 4.14 | 31,271 |
| Rep2-MazF | MazF | 21.02 | 15.59 | 2.96 | 30,338 |
| Rep2-FTO | FTO-MazF | 18.49 | 13.56 | 2.75 | 30,614 |
| Rep3-MazF | MazF | 23.62 | 17.67 | 3.21 | 30,490 |
| Rep3-FTO | FTO-MazF | 15.00 | 10.19 | 2.23 | 30,129 |

* Q20 indicated mapping quality >= 20.

**Table S2. Designed probes and universal primers for T3 ligase-based validation.** rG and rU indicates ribonucleotide.

| Chr. | Location | Probe | Sequences |
|---|---|---|---|
| chr4 | 6718671 | probe L | CCATAAGCAGGATTTGCAATTCACTTCATACTGGAGGTAAG ATATCGACCTGTACCTCT |
| chr4 | 6718671 | probe R | ATCTCATCCCTGCGTGTCAAGTAAACTTTAAAATCCCACACT CCAACATCATATATrGrU |
| chr9 | 119462465 | probe L | CACTGAAAGGTGGAGCAGATCCCACTGAAGCATGCACAGT GATATCGACCTGTACCTCT |
| chr9 | 119462465 | probe R | ATCTCATCCCTGCGTGTCACTGAGACATAATGTTACGGGAA TATGATGTCTTAAATrGrU |
| chr1 | 29070177 | probe L | CCACTGCCGTTGACACTGAAAAGTAAGTAAACGGGGCCTTG ATATCGACCTGTACCTCT |
| chr1 | 29070177 | probe R | ATCTCATCCCTGCGTGTCGTCCACAGCAGATTTCATTTCTGC CACGCCACAGAAGTrGrU |
| chr22 | 38202158 | probe L | CCGCAAGAAAAGACTTCATTGTCACTTCTTCTTGCCGGCCG ATATCGACCTGTACCTCT |
| chr22 | 38202158 | probe R | ATCTCATCCCTGCGTGTCAGAAAATTATTTACAGAAAATAG GAGACAGGAGGGAGTrGrU |
| chr16 | 69362741 | probe L | CCGTAACTGATTTCAGGGCAAACATTTCTGACATCTTCCTGA TATCGACCTGTACCTCT |
| chr16 | 69362741 | probe R | ATCTCATCCCTGCGTGTCGGGGGTGATTTTTCTCCTCAAGTT GTAGCCAACATTTTrGrU |
| chr22 | 21307376 | probe L | TCATATTCCTCTGAGCAAACATCACAAGAGCAGTGGCCATG ATATCGACCTGTACCTCT |
| chr22 | 21307376 | probe R | ATCTCATCCCTGCGTGTCTGGCAAAAAACCACATCATCCCTT TTCAAAAATAAAATrGrU |
| chr1 | 227152794 | probe L | GGAGGCTCCGGCTGCATGCGGGACTGAGAAGTGGAACTCC GATATCGACCTGTACCTCT |
| chr1 | 227152794 | probe R | ATCTCATCCCTGCGTGTCGCGCTGACTGGTCGGGAGCGGAG GCTGAAGAGAAGTCTrGrU |
| chr3 | 99886615 | probe L | GGTTCTTCCTAAGTTTTACCCACTTAGGACAAATTTCTTTGA TATCGACCTGTACCTCT |
| chr3 | 99886615 | probe R | ATCTCATCCCTGCGTGTCCTGCAGATGATCAGCATCAGGAC CGATTTCTTCTCACTrGrU |
| chr19 | 41785018 | probe L | AGACGGAAGGTGCTTCACGGAGATTTCCTCATTGATCTTCG ATATCGACCTGTACCTCT |
| chr19 | 41785018 | probe R | ATCTCATCCCTGCGTGTCGGGACCAGCCGATACGGACCACG TGGGGGTCAGGCTCTrGrU |
| Universal primers | | FP | CCATCTCATCCCTGCGTGTC |
| Universal primers | | RP | AGAGGTACAGGTCGATATCA |

**Table S3. Designed primers for quantitative PCR validation.**

| Chr. | Location | Probe ID | Sequences |
|---|---|---|---|
| chr16 | 87865500 | H1-FP | TGTCACGCTCATGTCCTGTC |
| chr16 | 87865500 | H1-RP | CACGGGAACAACAGAAACAA |
| chr6 | 31130366 | H2-FP | TCCTCGGAAGTACCCAGTGA |
| chr6 | 31130366 | H2-RP | ACATGGAACCAGACGTCACA |
| chr19 | 41785018 | H3-FP | TTGCATACCTGTGGTCAGGA |
| chr19 | 41785018 | H3-RP | AGGAGAAAGGCTCTTCGCCT |
| chr9 | 131763970 | L1-FP | TCCTCACCACTCTAGAGGTG |
| chr9 | 131763970 | L1-RP | GTTGGTGCTCAGCTGGTACA |
| chr19 | 59059888 | L2-FP | TCAATGCCTGGACCAAGAGT |
| chr19 | 59059888 | L2-RP | ATAGCCTTCCTGCACCTCCA |
| chr16 | 30537003 | L3-FP | TCAACTGGTGTCTGGCTGAT |
| chr16 | 30537003 | L3-RP | CACCATAGGAGAGTCTCAGT |
| -- | -- | Gapdh-FP | TGAGTACGTCGTGGAGTCCA |
| -- | -- | Gapdh-RP | TTCACACCCATGACGAACAT |

**Table S4. Basic information of sequencing data for mammalian tissues.**

| Species | Tissue | Treatment | Clean data (million reads) | Mapped reads (million reads) | Mean coverage (Q20*) | # covered transcripts |
|---|---|---|---|---|---|---|
| Human | Brain | MazF | 24.62 | 18.19 | 2.20 | 35,285 |
| Human | Brain | FTO-MazF | 24.79 | 18.24 | 2.11 | 35,295 |
| Human | Liver | MazF | 26.55 | 19.44 | 2.43 | 33,849 |
| Human | Liver | FTO-MazF | 27.61 | 19.92 | 2.24 | 33,695 |
| Human | Kidney | MazF | 26.58 | 18.36 | 2.30 | 34,859 |
| Human | Kidney | FTO-MazF | 23.03 | 16.48 | 2.03 | 34,660 |
| Mouse | Brain | MazF | 14.13 | 7.46 | 1.82 | 46,194 |
| Mouse | Brain | FTO-MazF | 29.43 | 17.71 | 2.52 | 47,537 |
| Mouse | Liver | MazF | 21.53 | 11.73 | 2.82 | 42,522 |
| Mouse | Liver | FTO-MazF | 21.05 | 11.45 | 2.75 | 42,194 |
| Mouse | Kidney | MazF | 18.71 | 10.99 | 2.17 | 44,831 |
| Mouse | Kidney | FTO-MazF | 27.30 | 15.57 | 2.48 | 45,584 |
| Mouse | Heart | MazF | 15.76 | 8.31 | 1.96 | 44,363 |
| Mouse | Heart | FTO-MazF | 49.02 | 27.23 | 6.88 | 46,070 |
| Mouse | Testis | MazF | 17.09 | 10.41 | 1.93 | 52,981 |
| Mouse | Testis | FTO-MazF | 25.47 | 14.83 | 2.23 | 53,674 |
| Rat | Brain | MazF | 10.90 | 7.26 | 1.58 | 54,593 |
| Rat | Brain | FTO-MazF | 12.81 | 8.14 | 1.60 | 55,004 |
| Rat | Liver | MazF | 12.55 | 7.52 | 2.31 | 48,170 |
| Rat | Liver | FTO-MazF | 15.82 | 9.75 | 2.58 | 48,489 |
| Rat | Kidney | MazF | 14.83 | 9.13 | 2.06 | 53,011 |
| Rat | Kidney | FTO-MazF | 14.45 | 8.67 | 1.93 | 52,712 |

* Q20 indicated mapping quality >= 20.

**Table S5. m$^6$A sites identified by m$^6$A-REF-seq for mammalian tissues.**

| **Species** | **Tissue** | **# m$^6$A sites** |
|---|---|---|
| Human | Brain | 9,244 |
| Human | Liver | 5,312 |
| Human | Kidney | 9,225 |
| Mouse | Brain | 16,161 |
| Mouse | Liver | 8,318 |
| Mouse | Kidney | 7,938 |
| Mouse | Heart | 4,419 |
| Mouse | Testis | 9,456 |
| Rat | Brain | 4,720 |
| Rat | Liver | 3,554 |
| Rat | Kidney | 6,907 |

**Table S6. Shared m$^6$A-modified genes and m$^6$A sites between pairwise species and significance.** The hypergeometric test was used to test the significance.

| Brain | shared-m$^6$A genes | | | Brain | shared-m$^6$A sites | | |
|---|---|---|---|---|---|---|---|
| | Human | Mouse | Rat | | Human | Mouse | Rat |
| Human | | 1,456 | 704 | Human | | 229 | 105 |
| Mouse | $8.6\times10^{-311}$ | | 1,003 | Mouse | $1.2\times10^{-26}$ | | 267 |
| Rat | $9.8\times10^{-197}$ | $2.7\times10^{-322}$ | | Rat | $2.1\times10^{-30}$ | $1.2\times10^{-105}$ | |

| Kidney | shared-m$^6$A genes | | | Kidney | shared-m$^6$A sites | | |
|---|---|---|---|---|---|---|---|
| | Human | Mouse | Rat | | Human | Mouse | Rat |
| Human | | 1,065 | 911 | Human | | 136 | 108 |
| Mouse | $3.2\times10^{-228}$ | | 947 | Mouse | $5.5\times10^{-20}$ | | 190 |
| Rat | $5.0\times10^{-212}$ | $1.4\times10^{-256}$ | | Rat | $5.0\times10^{-17}$ | $7.5\times10^{-68}$ | |

| Liver | shared-m$^6$A genes | | | Liver | shared-m$^6$A sites | | |
|---|---|---|---|---|---|---|---|
| | Human | Mouse | Rat | | Human | Mouse | Rat |
| Human | | 664 | 395 | Human | | 97 | 65 |
| Mouse | $2.2\times10^{-154}$ | | 609 | Mouse | $4.0\times10^{-23}$ | | 158 |
| Rat | $4.6\times10^{-139}$ | $2.1\times10^{-213}$ | | Rat | $2.3\times10^{-28}$ | $3.1\times10^{-80}$ | |

### Supplementary Figures

**Figure S1. Quantitative demonstration of various fractions of m⁶A-containing oligo mixed with unmethylated oligo digested by ChpBK.**

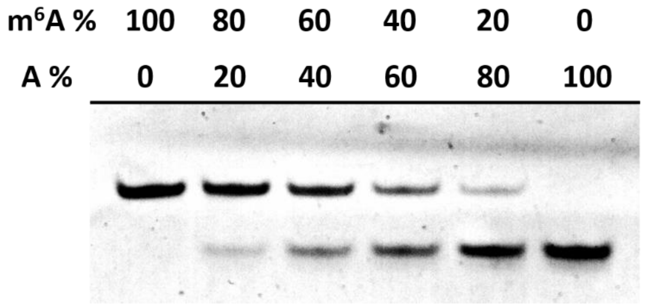


**Figure S2. FTO demethylate assay.** (a) Dot-blot assay. (b) Detection of FTO demethylation reaction with different FTO concentration by MazF cleavage assay.

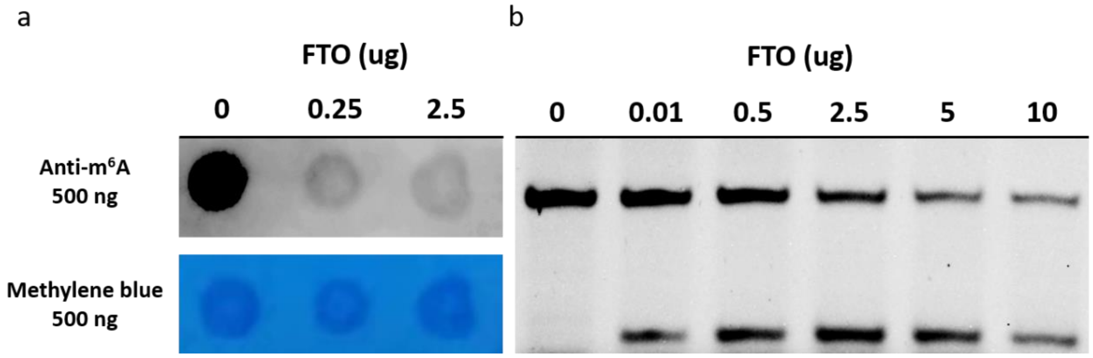

**Figure S3. The base composition of mRNA products cleaved by MazF.** The x-axis indicates the relative position to 5' terminal. Most reads contained the ACA 5' end as shown in the position 1, 2 and 3.

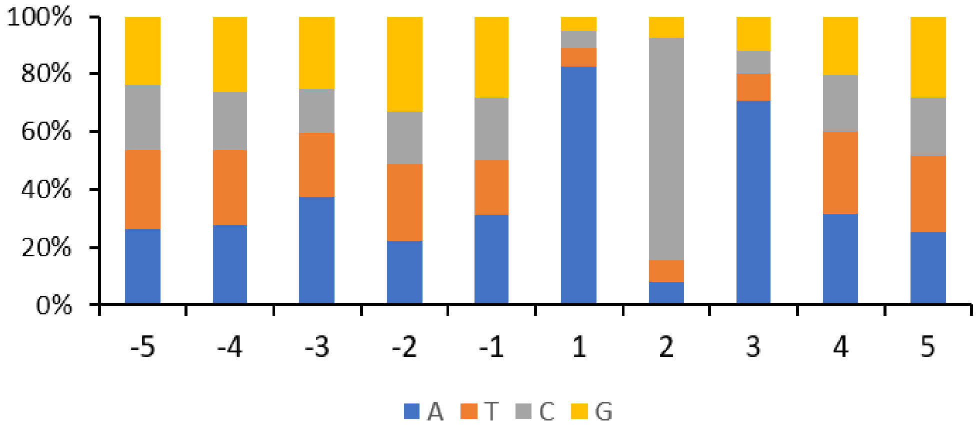


**Figure S4. The single-site validation for 18S rRNA control site and m$^6$A site.** The ligation reaction with 500 ng total RNA template was performed at room temperature for 10 min. Different amounts (1/10 ul to 1/50 ul) of T3 ligase was used. FTO- indicated the reaction without FTO demethylation while FTO+ indicated the FTO treatment. The 1/50 ul T3 ligase reaction was able to distinguish the m$^6$A from A. M indicated the 50 bp marker, ranging from 50 bp to 600 bp.

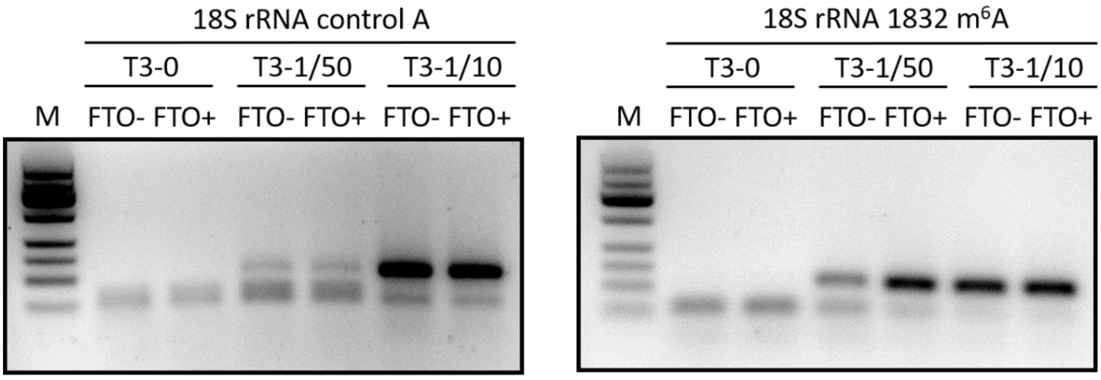

**Figure S5. The single-site validation of eight m$^6$A sites and one unmethylated site.** Eight m$^6$A sites and one A site (chr22: 21307376) were picked from m$^6$A-REF-seq. Six out of eight m$^6$A sites were confirmed to be m$^6$A sites. The amount of T3 ligase (ul) and ligation time in each reaction were indicated in the figures. FTO- indicated the mRNA without FTO demethylation while FTO+ indicated the FTO treatment. M indicated the 50 bp marker, ranging from 50 bp to 600 bp.

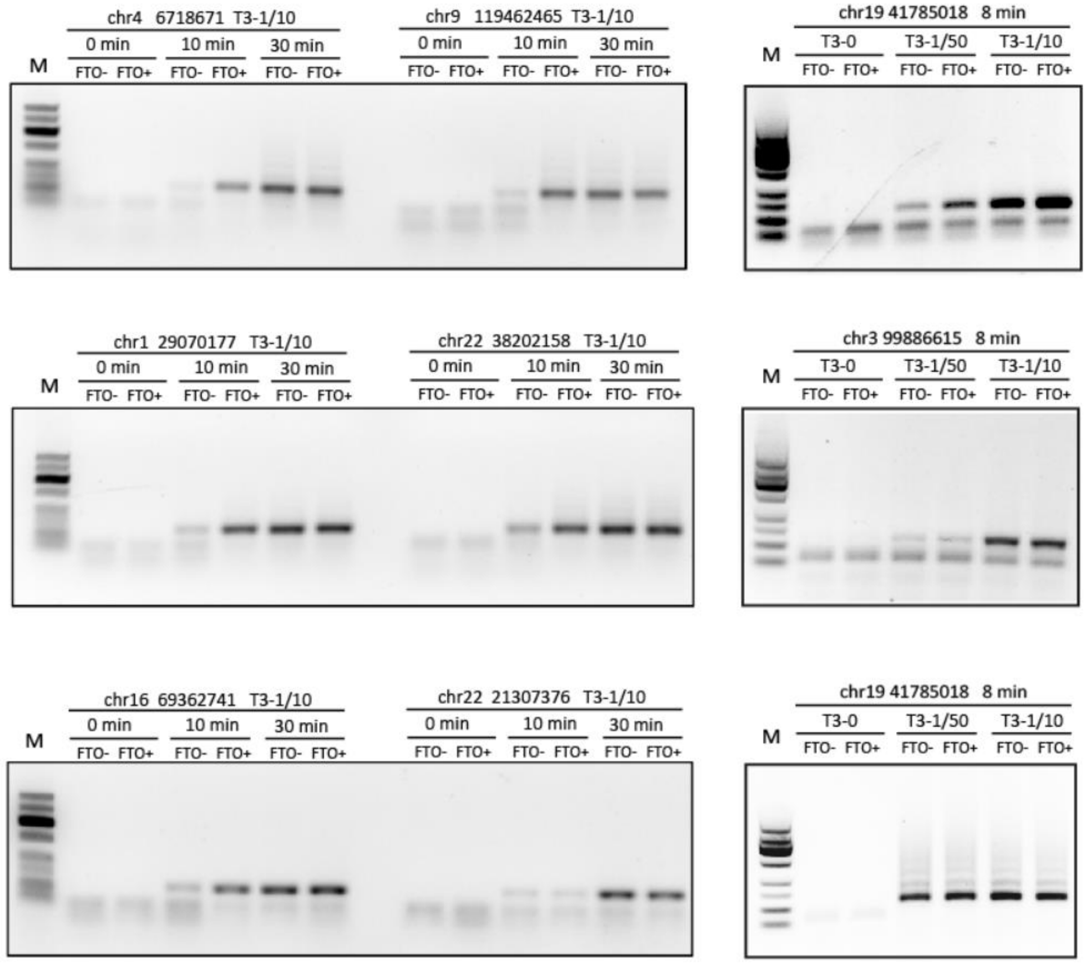

**Figure S6. Quantitative PCR results of six m$^6$A sites.** H1-H3 were high level methylated sites (> 0.75) while L1-L3 were low level methylated sites (< 0.35). See Supplementary Table S3 for designed primers.

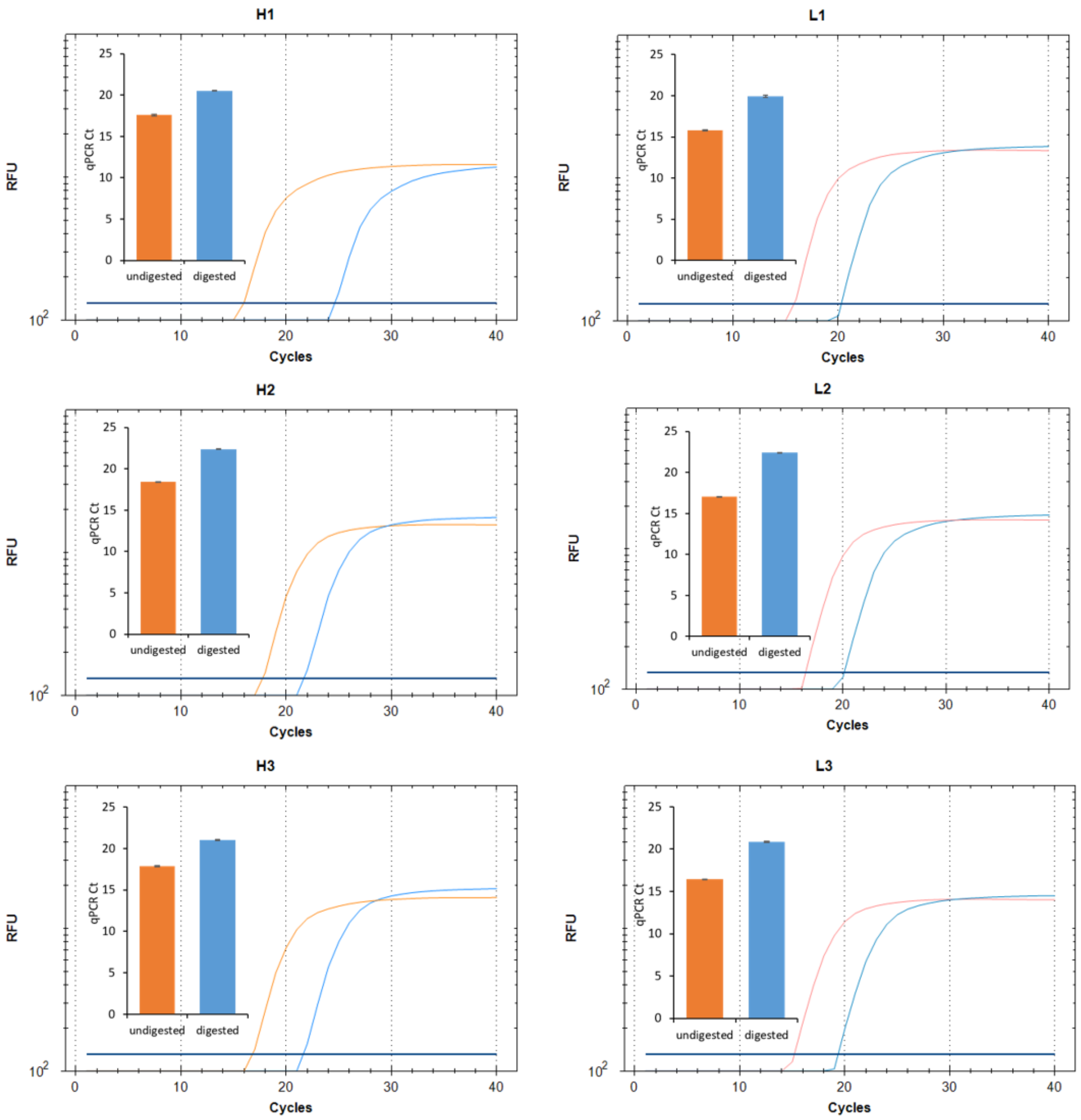

**Figure S7. Metagene plots of m$^6$A in mouse heart and mouse testis.**

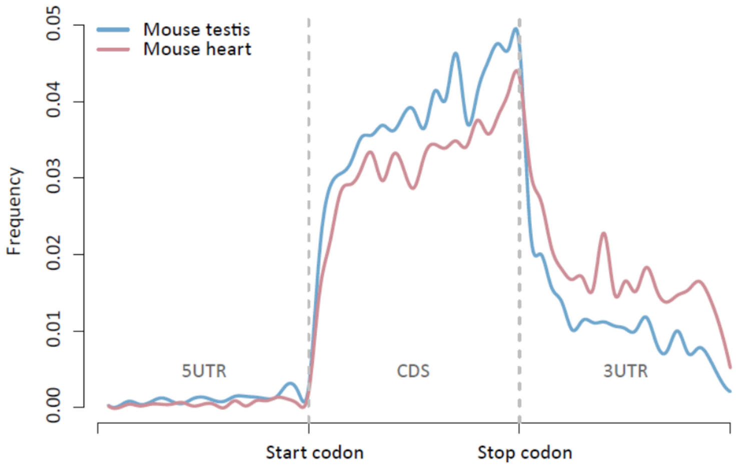

**Figure S8. Conservation of m$^6$A in mammalian kidney.** (a) Metagene plots of m$^6$A in the kidney of human, mouse and rat. (b) Shared m$^6$A-modified genes among three species. (c) Diagram showing the m$^6$A sites conserved in the corresponding short regions from different species. (d) Frequency of distances for pairwise m$^6$A in kidney. Randomly picked ACA motifs were assigned for the same analysis as control. (e-f) Conservation scores of all m$^6$A sites, methylation sites in ortholog genes and conserved m$^6$A sites were compared to that for all A sites in ACA motifs (Wilcoxon test, p-values < $2\times10^{-7}$).

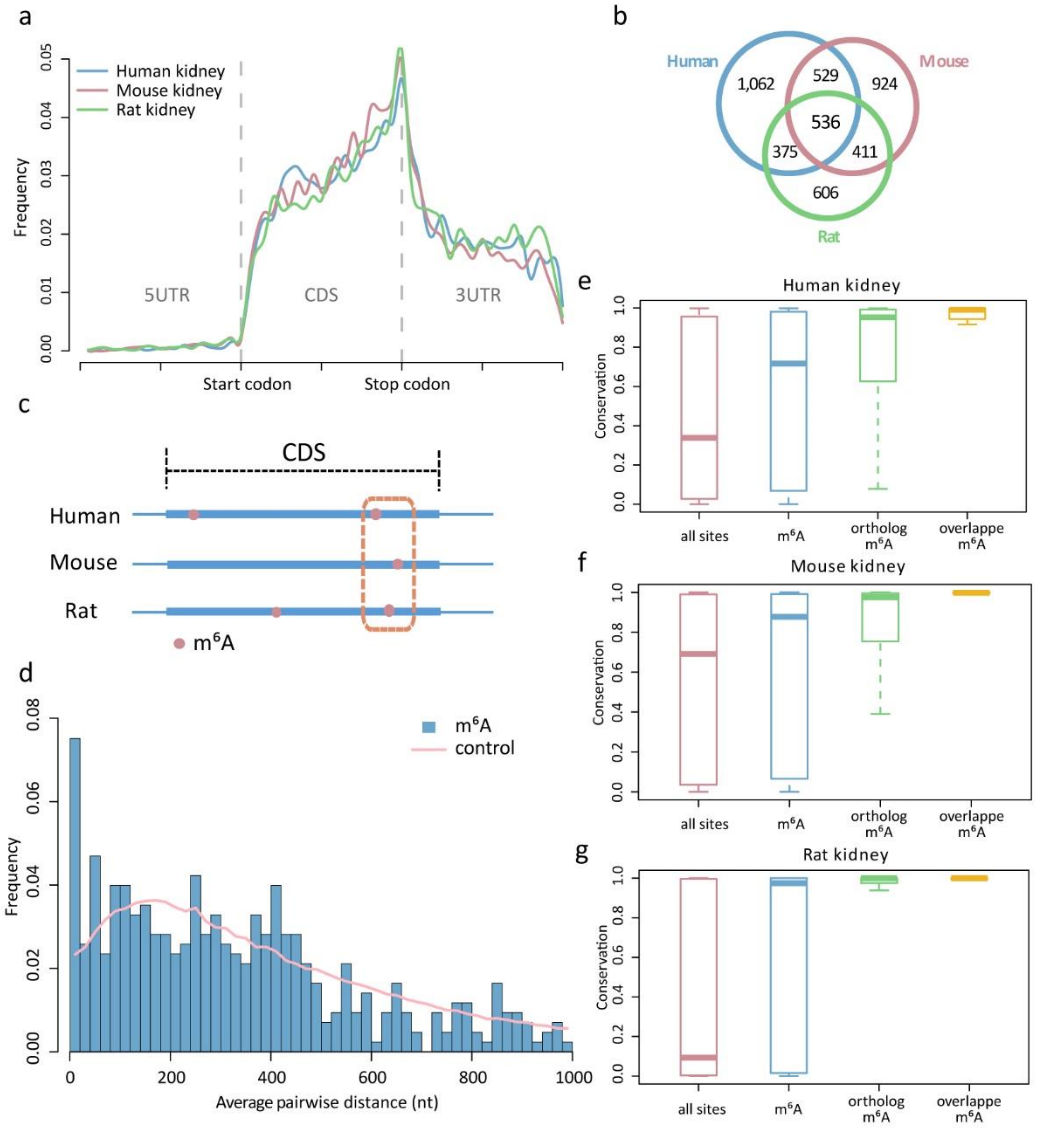

**Figure S9. Conservation of m$^6$A in mammalian liver.** (a) Metagene plots of m$^6$A in the liver of human, mouse and rat. (b) Shared m$^6$A-modified genes among three species. (c) Diagram showing the m$^6$A sites conserved in the corresponding short regions from different species. (d) Frequency of distances for pairwise m$^6$A in liver. Randomly picked ACA motifs were assigned for the same analysis as control. (e-f) Conservation scores of all m$^6$A sites, methylation sites in ortholog genes and conserved m$^6$A sites were compared to that for all A sites in ACA motifs (Wilcoxon test, p-values < 0.0003).

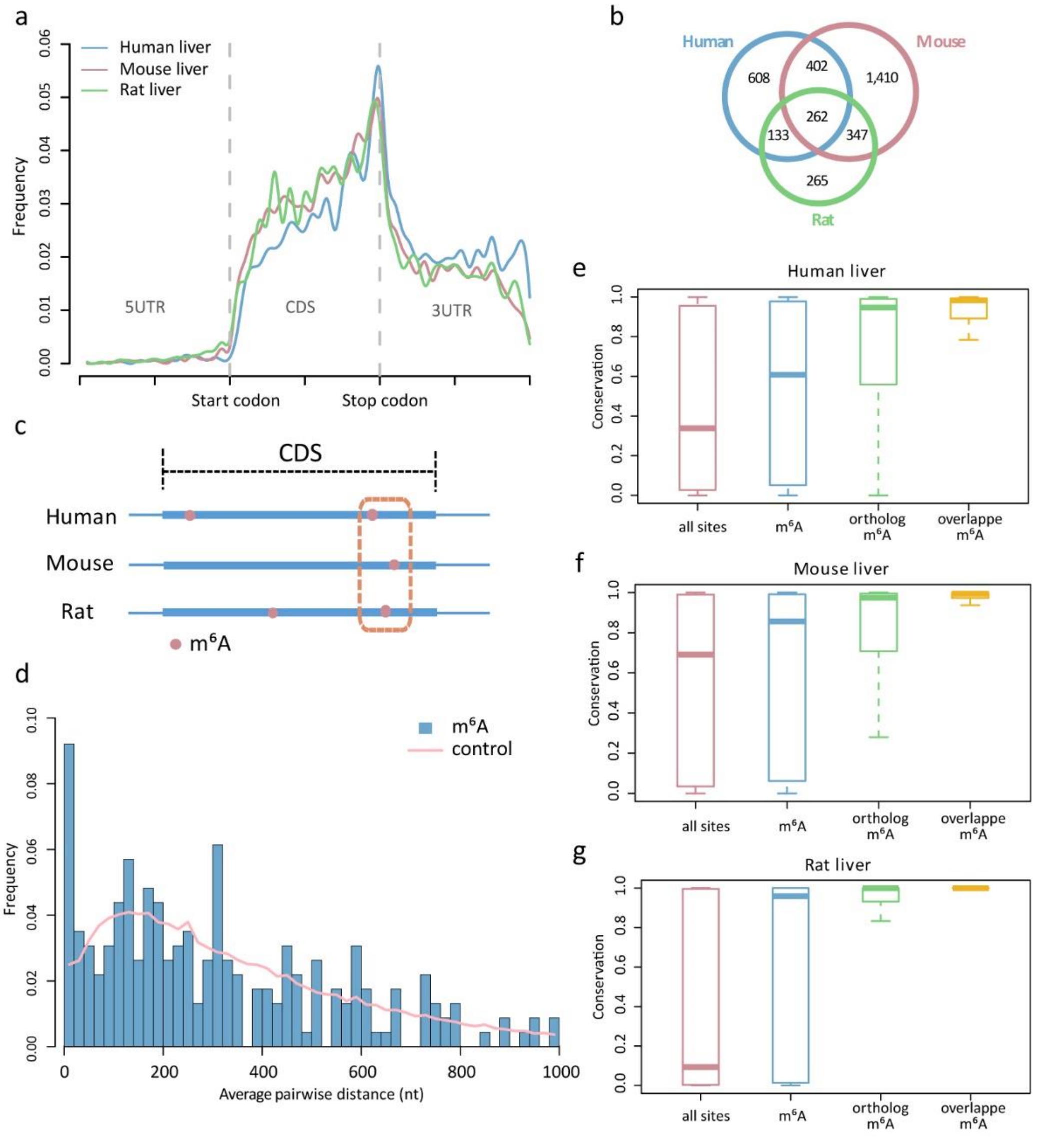

**Figure S10. Conservation scores for brain.** Conservation scores of all m$^6$A sites, methylation sites in ortholog genes and conserved m$^6$A sites were compared to that for all A sites in ACA motifs (Wilcoxon test, p-values $< 9\times10^{-10}$).

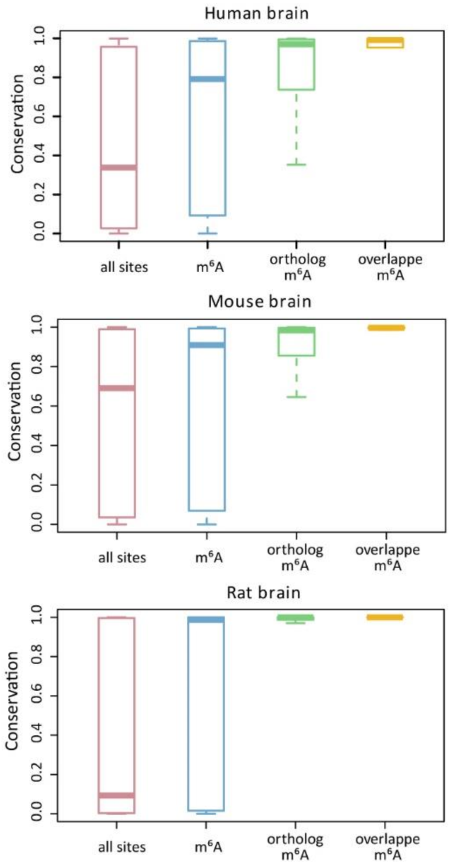


**Figure S11. Validation for the methylation sensitivity of mutated MazF-K56A.** Two synthetic RNA oligonucleotides were used as the substrates (RB1 and RNA). The sequences can be found in Methods section.

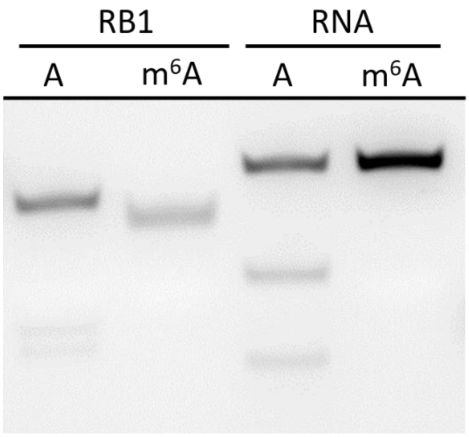
